# Divergent pattern of development in rats and humans

**DOI:** 10.1101/2023.04.11.536227

**Authors:** Wanda campos, Tomas Iorri, Antonella Presti, Rafael Grimson, Pablo Vázquez-Borsetti

## Abstract

*Rattus norvegicus* is the second most used laboratory species and the most widely used model in neuroscience. Nonetheless, there is still no agreement regarding the temporal relationship of development between humans and rats. We addressed this question by examining the time required to reach a set of homologous developmental milestones in both species. With this purpose, a database was generated with data collected through a bibliographic survey. This database was in turn compared with other databases about the same topic present in the literature. Finally, the databases were combined, covering for the first time the entire development of the rat including the prenatal, perinatal, and postnatal periods. This combined database includes 497 individual dates for both species. The developmental relationship between humans and rats was better fit by a logarithmic function than by a linear function. Also, an inflection point close to birth becomes evident after the logarithmic transformation of the data. The predictions of the proposed model were compared with other estimators historically used to calculate developmental relationships, such as growth in brain weight, finding a notorious discrepancy. As development progresses, an increase in the dispersion of the data is observed. Until now, developmental relationships have been interpreted as a univocal equivalence. In this work, is proposed an alternative interpretation where the age of one species is translated into a range of ages in the other.

## Introduction

The use of animal models is essential for scientific development in the fields of biology and medicine. Their importance not only involves the generation of new knowledge. It is also indispensable for translating basic scientific findings into therapeutic interventions for patients. Success in this task depends on the selection of an appropriate model. At the same time, the similarities of these species with humans are always in question. This becomes even more complex in the case of development. Rattus norvegicus is the second most used laboratory species and the most widely used model in neuroscience but there is still no agreement regarding the temporal relationship of development between humans and rats (Ortiz et al., 2022). Translating ages across species has proven to be a difficult task, and the case of neurodevelopment represents an even bigger challenge due to its complexity. Misinterpretation of these relationships can lead to errors that increase the difficulty of extrapolating results to humans, waste scientific resources and pose an ethical problem regarding the inefficient use of laboratory animals.

Although there are many works that have addressed this problem with different degrees of depth(Clancy et al., 2001, 2007; Donaldson, 1918; Dutta & Sengupta, 2016; Finlay, 2008; Hamdy et al., 2020; Romijn et al., 1991; Semple et al., 2013; Vargas Mamani, 2020; Workman et al., 2013) there is still no agreement regarding the mathematical model that better explains the developmental relationships between both species or how it should be interpreted. Most of these works proposed a linear relationship. On the other side, Dr. Barbara Clancy (Clancy et al., 2001, 2007; Workman et al., 2013) proposed a logarithmic function to model the developmental equivalence of a handful of species including the rat. That work proposed a general rule that describes relationships in the development of different species, but it was not intended to specifically compare humans and rats. More recently, Ohmura and Kuniyoshi (Ohmura & Kuniyoshi, 2017) conducted a study specificaly for the rat and questioned the results obtained by Clancy. They presented an open database with more data and proposed that the developmental relationships could be explained with a linear relationship if birth was not taken into consideration as a developmental milestone. The authors decided to treat the birth as an “outlier” and exclude it from the analysis.

At the end, both Clancy and Ohmura coincided in designating the birth as an “outlandish” milestone of development.

In this work we will analyze the temporal pattern of developmental milestones in rats and humans, paying special attention to neurological development. A logarithmic model will be proposed to explain the relationship between the timeline of human development, expressed in days after fertilization (DAF), and the DAF of rats as dependent variable. We will also criticize the canonical interpretation of a univocal relationship in favor of a comprehensive relationship that translates ages into ranges of time.

## Methods

A literature survey was conducted to collect data and public databases with relevant information about the comparative development of humans and rats. Three databases comparing the age of both species were found (Clancy et al., 2001; Godlewski et al., 1997; Ohmura & Kuniyoshi, 2017). Also, a fourth database was created specifically for this work. The data was compiled from search engines such as National Center for Biotechnology information (NCBI)/PubMed, Google Scholar, ResearchGate and Elsevier/ScienceDirect without restriction of years. Google drive and Excel spreadsheets from the Microsoft Office package were used to compose the database. In addition, the Mendeley application was used as a citation manager.

From now on, each database will be referred to by the name of the author, being the one made for this paper called Campos-Iorii.

All the data reported in this work was obtained entirely from original empirical works present in the literature. In a preliminary analysis, a substantial amount of information from reviews and books that could not be traced back to their original sources was found. Although in many cases revisions of literature were used to find some of the research papers mentioned above, information that could not be traced to the original source was not included.

Ohmura’s database was created with a similar intention than this work. The author proposed 94 specific points of data for both species, of which 20 were perinatal milestones. This database included some milestones of mouse development (16) that had to be removed to improve the specificity of the comparison with rats.

Clancy’s database proposed a final number of 11 paired data points for rats and humans. Also were included a comparative morphological study of the first days of development carried out by Godlewski with 13 data points from the first embryonic stages.

All the databases were reformatted to the structure explained in the following section.

### Database structure

The database was structured in rows that correspond to developmental milestones or processes of one of the two species. Campos-Iorii database contains 293 rows with 143 rows of data corresponding to rats and 150 corresponding to humans. There are three types of data:

Punctual: It describes a singular development milestone, there are 76 for each species, corresponding to 152 rows of the database created for this work, to which must be added 26 data rows from Godlewski’s database, 22 from Clancy’s database and 156 from Ohmura’s database.

Range: It is a subgroup of punctual data, where the beginning and the end of a process is reported, 8 of these processes were included in this work. This type of data occupies 64 rows and is only present in the database generated for this work.

Contiguous data: includes data that were obtained from works where the progress was studied in a sequential and quantitative fashion, i.e., the weight of the brain through development. In order to compare both species, the values were normalized to the adult average, that was considered 100%. This type of data was not included in the generation of the models. Three phenomena were analyzed: increase in brain weight, increase in GAD (glutamate decarboxylase a GABA-producing enzyme) activity in the cortex, and increase in GAD activity in the cerebellum. These kinds of data occupy 124 lines in the database and are only present in the database generated for this work.

The combination of the 4 databases ends up generating a database with 497 rows of information for both species.

Each row contains the following columns:

Species: Indicates the correspondence with the species, Rat or Human.

Parameter: the neurodevelopmental milestone or the process to be compared, i.e. Establishment of the blood-brain barrier.

Value: meant to report quantitative data. If the data is not quantitative, the placeholder “Punctual” was annotated for punctual data. Also, if the data corresponds to a range the “Start” and “End” of the process was annotated in this column. It may be important to emphasize that the three types of data are associated in other columns with the corresponding ages.

Pcdays: age in postconceptional days. Postnatal data in the bibliography is usually related to postnatal age, for this reason were added the intrauterine age of 22 days for rats and 280 days for humans. The bibliography differs in the mechanism used to determine the moment of fertilization. In most cases, especially in humans, it approximates the ovulation date, generating an error of 2 weeks for humans and 1.5 days for rats. Unless otherwise stated in the publication, embryonic/fetal days, gestational days, and post-conception days were considered equivalent. The day of birth was considered as postnatal day 0 (PND0). The data for years was approximated to days by multiplying by 365.25 and the data for months by 30.43.

PCGP: is a mathematical relationship between the age at which the milestone occurs and the gestation period of the species. Useful to quickly assess the data, input errors and outliers.

%: in the case of quantitative contiguous data, the percentage is indicated here, taking the average adult value as 100%.

Citation: the source from which the data was obtained in abbreviated form.

Obs.: information relevant and/or complementary to the data was mentioned there.

Quote: this column is intended to simplify the search in the original source of the data, showing the textual citation, figure, or table in which, it was mentioned.

DOI: the Digital Object Identifier (or DOI), a unique code of each cited publication, was included there.

### Statistical analysis

The programs for data analysis and graph generation were developed in-house using the Python programming language (Python 3.12, Wilmington, DE: Python Software Foundation). All the libraries (NumPy, Pandas, Scipy, Statsmodels, Scikit-learn, Matplotlib and Seaborn) and the IDE (Integrated Development Environment) Spyder were obtained through the Anaconda distribution. Because the coefficient of determination is not the best estimator to evaluate nonlinear data, we decided to utilize the Bayesian Information Criterion (BIC) and/or the Akaike Information Criterion (AIC). These tests perform better than the coefficient of determination with nonlinear models (Spiess & Neumeyer, 2010). The distribution of the data also presents high heteroscedasticity making the coefficient of determination even less appropriate to compare models. The statsmodels library provides important information about the models including BIC and AIC, and both were used to assess the models. In the supplementary materials were included the summaries of information for the linear regression and nonlinear regression.

The confidence intervals of the model parameters were analyzed by bootstrapping.

Both the database and the code generated for the analysis are available in the supplementary materials and through the link on the Github website reserved for the publication of the database:

https://github.com/Vazquez-Borsetti/rat-and-human-comparative-development

The last row shows the unification of all the previous databases into a single database (e).

## Results

### The relationship of development between humans to rats fits well to a logarithmic model

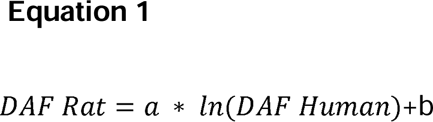

DAF= Days After Fertilization

To assess the developmental relationships between rats and humans, the temporal distribution of different events in the development of both species was analyzed. The independent variable corresponds to the human age in DAF and the dependent variable is the rat age in DAF. Then, a pair of coordinates can be assigned to each specific developmental event present in the database. The two competing mathematical models proposed in the bibliography to describe this relationship (linear and logarithmic) were tested. Nonlinear and linear ordinary least squares regressions were utilized to analyze the results. The temporal relationship of rat and human developmental milestones fits well to a logarithmic function. In other words, when we compare the development of the human with the rat, it is observed that the relative distance between development milestones increases with the age of the human. Figure 1 shows the database generated for this work as well as other aforementioned studies. These studies cover different stages of development. All of them fit well to a logarithmic function adjusted by ordinary least squares. The logarithmic fit outperformed the linear fit in our database (Campos-Iorri) and in all datasets combined together. The other databases that covered short periods of time (Godlewski, Clancy, Ohmura) presented a better fit with the linear regression. In these last cases the differences in the AIC and BIC are not substantial (see supplementary files statsnl.txt for the fit of the equation 1 and statsolslr.txt for the fit of the linear model). This can be expected because a linear regression can fit a non-linear curve at short intervals, the relationship may appear linear and consistent within a specific range. However, with larger intervals as in the case of our database and the combination of all databases, the non-linearity of the curve becomes more evident. **Additionally, when** the interception parameter was included, the linear fits of different databases showed different slopes and intercepted the ordinate axis at higher values as the database included later developmental milestones. This does not make any theoretical sense. The opposite can be observed with the logarithmic fit, where the parameters tend to be more consistent and approximate better to 0. In the supplementary materials are included two graphs showing the residuals for the linear (supplementary figure 1) and logarithmic (supplementary figure 2) models for each database. It becomes evident that a linear regression cannot explain late developmental events and that there is no consistency among different databases when this model is used. The opposite can be observed in the logarithmic model.

**Figure 1:**
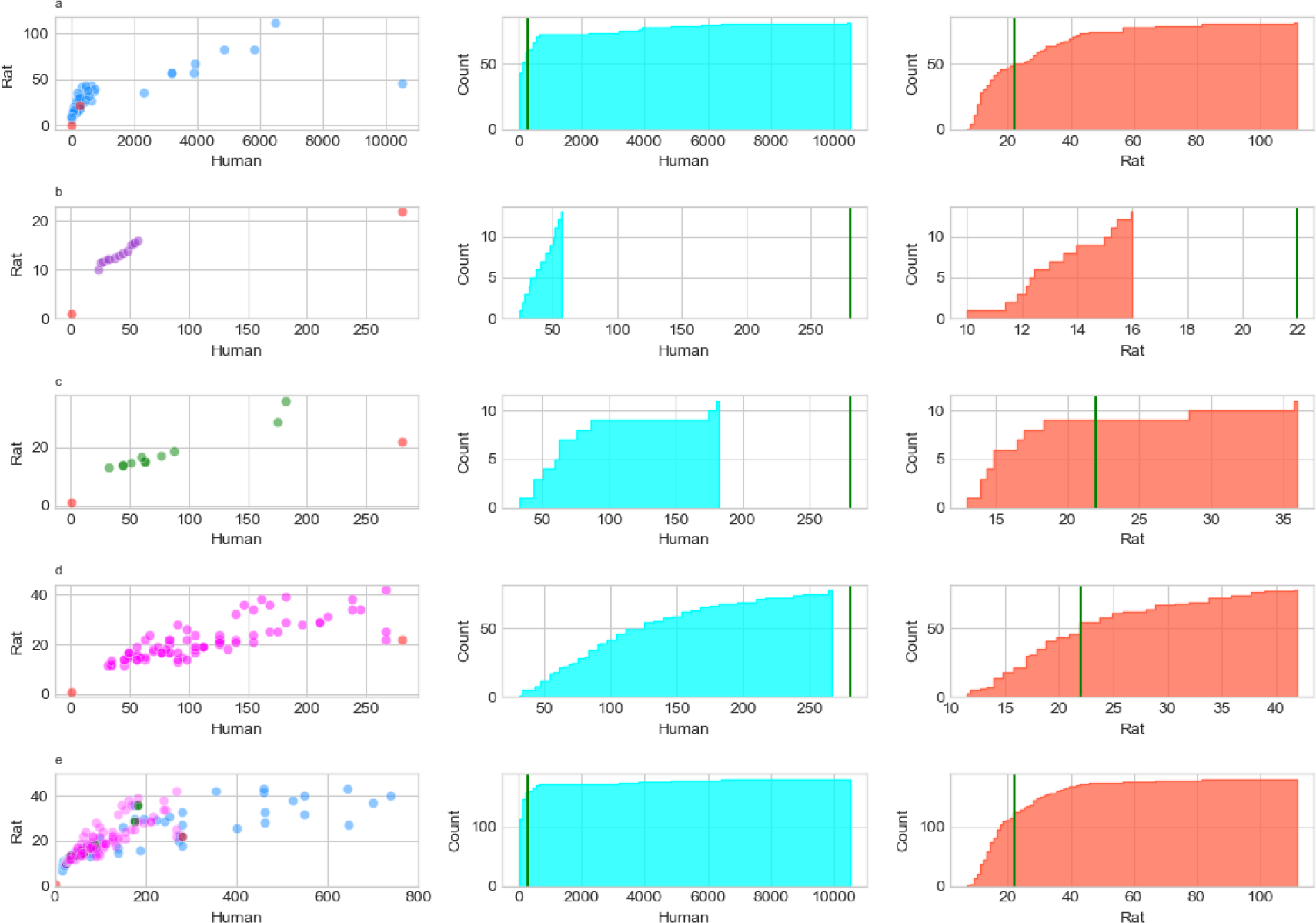
The first column shows the relationship of human and rat development (each dot represents a milestone in development). The axes are expressed in days after fertilization (DAF). The red dots indicate the dates of fertilization and birth. The second and third columns are cumulative histograms for human and rat data respectively. The green line marks the birth date. Each row corresponds to a database: Row 1 corresponds to the database made for this work (a), row 2 corresponds to the data from Godlewski’s comparative study(b), row 3 corresponds to Clancy’s database (c) and in the row 4 to Ohmura’s database (d).

**Figure 2:**
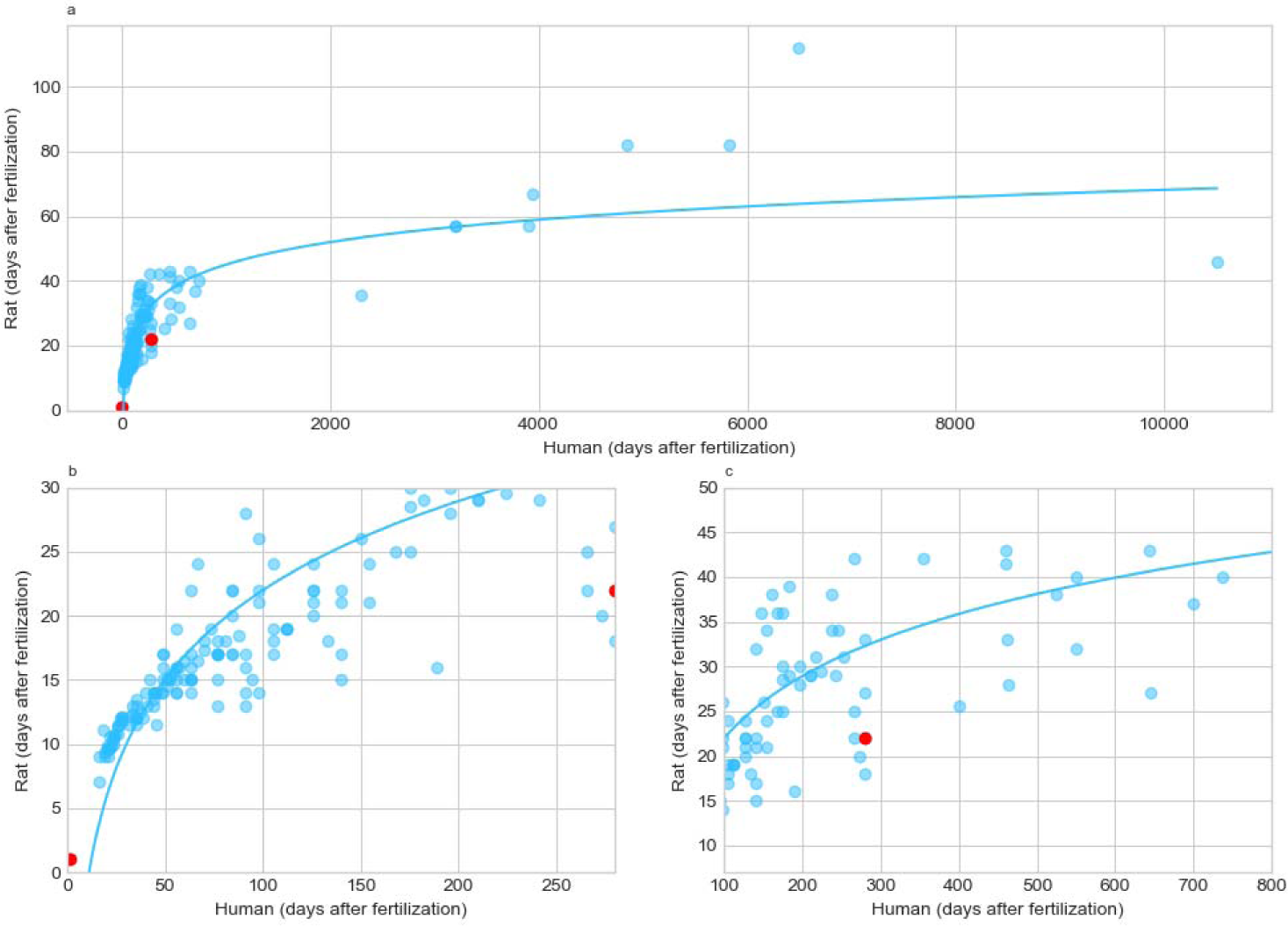
The first row shows ordinary least squares nonlinear regression (best fitting curve for equation 1) from the pooled database (a), the bottom rows show magnifications before (b) and around birth (red dot) (c).

In the residuals of the logarithmic model, also can be observed a slight change in the residual around birth. It is possible that birth changes the pattern of comparative development among both species introducing an inflection point in the distribution. Finally, all databases were combined into one (Figure 2) in order to obtain a mathematical model that describes the neurodevelopmental relationship between humans and rats (Equation 1: parameters: a=10.007, 95% CI [8.199, 11.897]; b= −24.098, 95% CI [−32.141, −16.5380]; CI were obtained by bootstrapping see supplemental image 5 and 6).

### The development of both species follows a divergent pattern

There is a complex relationship between the ages of both species. In our opinion, the canonical vision of a univocal relationship is an oversimplification. In figures 1 and 2 can be appreciated that the distribution of developmental milestones spreads out as the ages of the species analyzed increase. In other words, there is a divergent pattern of development in rats and humans. To analyze this pattern, a quantile regression was performed (figure 3). First, the data had to be linearized to facilitate the use of the linear quantile regression. Equation 1 was used with the parameters obtained in the non-linear regression to generate a new linearized independent axis that was called “human’”. This allows a more intuitive display of the data with a one-to-one relationship between developmental milestones for rats and humans.

**Figure 3:**
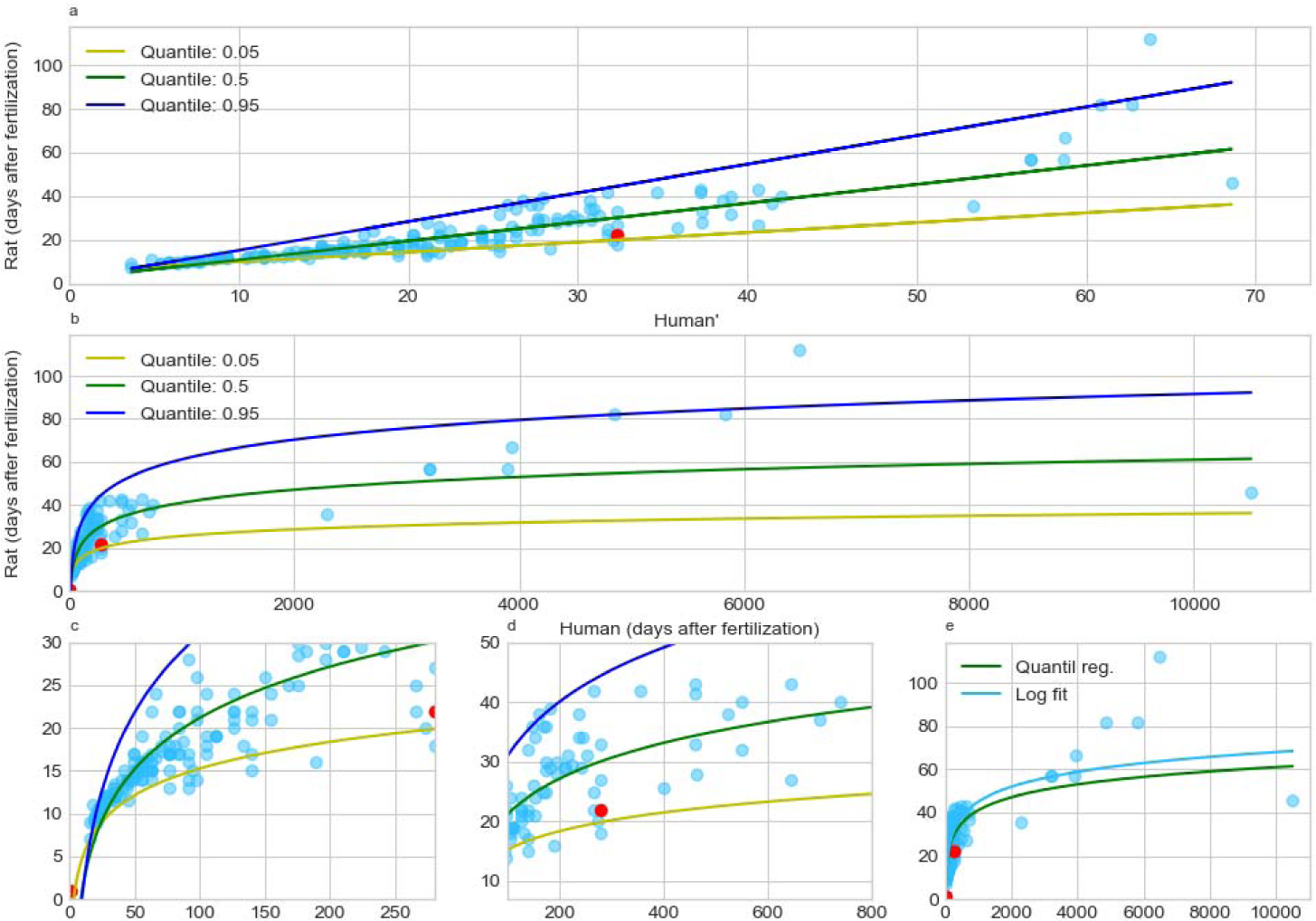
Quantile regressions of the pooled database. In the first row can be observed the linearization of the data using equation 1 with the parameters obtained in the ordinary least squares regression (a= 10; b= −24.1) and the subsequent generation of quantile regressions. The median quantile, that leaves 50% of the data below is represented with a green line; the upper quantile is the 95th *percentile* is represented and with a blue line; and the lower quantile is the 5th percentile *and* is represented with a yellow line. These last two quantiles comprehend 90% of the data. Note that the transformation with equation 1, in addition to linearizing the data, makes the scales comparable, for this reason the median quantile maintains a temporal relationship close to one (a). In the second row, the human age axis is restored, so the pattern recovers its logarithmic appearance (b). The third row shows an amplification of the events before birth and near birth which is marked with a red dot (c). The last figure in the third row shows the ordinary least squares nonlinear regression and the median quantile of the quantile regression (d).

The median quantile (50%) separates developmental events that are relatively delayed in the rat (above this quantile) from those that are relatively delayed in humans (below this quantile). The 95% and 5% quantiles generate a range where 90% percent of the data is comprehended. If we assume that our sample is representative of the population of all possible development milestones, this will mean that both margins delimit 90% of these events. The supplementary table 1 shows the equivalence of human age to rat age in ranges that enclose 90% of the developmental events. Both in the table and figure 3 can be observed that there is a large dispersion in the data and that it tends to increase with age.

Although there might have been measurement errors in the original studies, it is difficult to think that they could be responsible for the observed dispersion. There would be no reason for errors to increase as a function of age. It could be expected that these putative errors increase the number of outliers, but, as quantile regression is robust against outliers, it would not be expected to affect the model estimations. In other words, measurement errors shouldn’t noteworthy affect the limits estimated by this regression. Then, the increase in variability should be produced by differences in the development “schedules” of both species. According to our analysis, the human perinatal period would be equivalent (in a 90%) to a range that goes from 19 to 45 (22 gestational days +23 postnatal days) of development in the rat. Homologous developmental events present in a determined date of the development of one species span across a wide range of ages of the other species. Therefore, it makes no sense to translate ages as a univocal relationship. It is a better interpretation to consider these relationships as ranges. In summary, besides the fact that the distance between developmental milestones of humans increases with age in comparison with rats, as seen in the previous section, the observed pattern of development shows another characteristic: Different milestones and processes follow different time sequences. The relative times not only do not differ proportionally but also do not maintain the same temporal relationship. These two phenomena added together could explain by themselves both the variability and the observed increase in the dispersion of developmental milestones as age increases.

### Improving the linear approximation of ranges of developmental processes

In the preparation of the database, there were developmental milestones that included both the beginning and the end of a specific process i.e.: the start and the end of lactation. This data corresponds entirely to our database. A first attempt to establish the temporal equivalence of this process could be a simple linear approximation (Figure 4). However, as the relationship of development is not linear, a bias may be generated. To overcome this issue, the linearized axis “human’” was used as before (figure 4 b) to set the linear relationship and later recover the regular axis to make a better approximation. Therefore, if a process follows the same pattern in both species this method could allow us to estimate the degree of advance of the phenomenon in the other species. Unfortunately, this may not be common. As will be seen in the next section, different developmental processes do not necessarily follow similar temporal patterns in different species. Then this method should be more accurate than a linear approximation but may only be better for describing time relationships than other aspects relative to the process. To deepen into the complex relationships of comparative developmental processes the isometric development of two important processes will be described in the next section.

**Figure 4:**
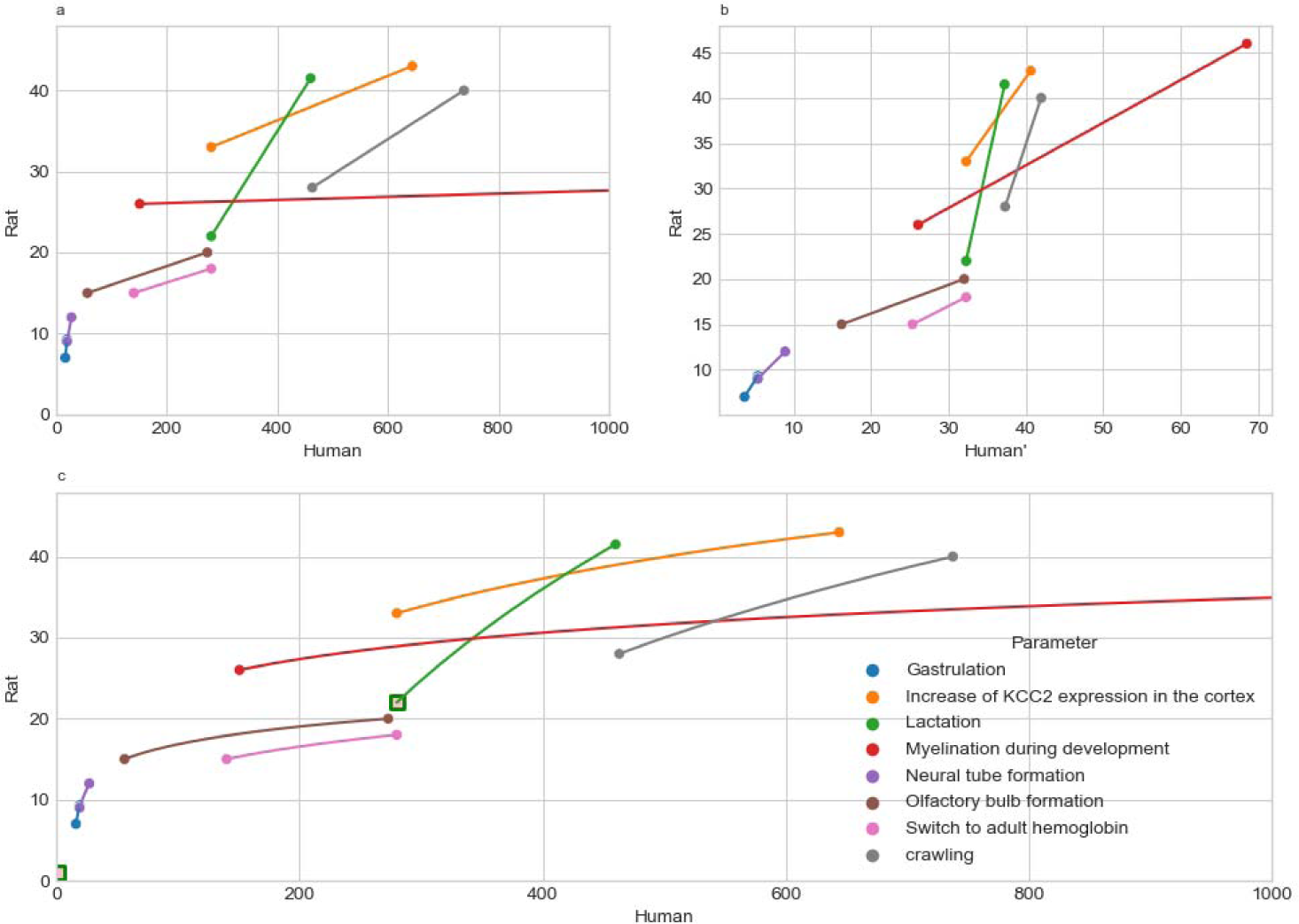
This figure shows several comparative development processes with known starting and ending points for both species. The lines that join the points would make it possible to estimate possible equivalences within the processes, but the fact that the age relationships are not linear generates a bias (a). In the second graph of the upper row, the relationship was linearized using equation 1 (b). In the second row, the third figure shows the temporal relationship when the original scale is recovered (c). These graphs give information about the temporal relationship of these processes but not necessarily about other equivalences. It is recommended to see the text for more detail in the interpretation of this graph.

### Brain weight is a moderate predictor of comparative brain development

There are processes such as the increase in brain weight that have been measured in detail in both species. After performing normalization and interpolation, it becomes possible to make isometric comparisons of this kind of data. In the case of brain weight, it becomes possible to know the corresponding date for each percentage of brain weight increase in humans and rats. These types of data were not considered to generate the models in the first and second section of this work. In consequence it is possible to compare these estimators of brain development against our model. The first comparison was the growth in brain weight to assess if it is a good estimator of the developmental relationships between both species. This is important because brain weight increase was historically used in the literature to estimate developmental parallels (Dobbing & Sands, 1979; Romijn et al., 1991). Also, the comparisons of developmental brain weight ratios between rats and humans are interesting by themselves. Human data about brain weight increase throughout development were taken from Dekaban and Sadowsky publications ((Dekaban & Sadowsky, 1978) which were subsequently reviewed by (Vannucci & Vannucci, 2019) 2019). Rat data were obtained from (Patterson et al., 2016; Weichenthal et al., 2010). To make the comparison, the data was normalized as a percentage of the mean adult value. The plots for humans and rats are shown in Figure5 (a;b). When there was a superposition of more than one measurement the values were averaged. The graph shows that the curve begins well below the median quantile and even below the level of 95% of events that develop more slowly in humans. At the time of birth, it is still below the median quantile. Approximately at the postnatal year of the human life and after 2 weeks postnatal life in the rat, the relationship between development and brain growth becomes slower in the rat. This occurs when around 60% brain weight gain is reached in both species. Finally, when the brain exceeds 80% of the adult average value, it also exceeds the upper limit of 95% of events that develop more slowly in the rat. Growth is not the same as development, it is at best a modest predictor, at least in the case of the nervous system. It is worth noting that the model includes 85% of the process, which is quite close to the expected 90%.

The last four subfigures plot continuous data of GAD enzyme activity in the cortex. The two graphs in the first row show the increase in enzyme activity as a percentage of the average adult value in relation to days after fertilization (DAF) throughout development for humans (e) and rats (f), respectively. The figures in the last row show the isometric relationship for the activity of the enzyme (g) and the overlap with our model (h).

### GAD activity is a bad predictor of comparative brain development

We also studied quantitative data on the activity of the GAD enzyme. The values are also relativized to the mean value for adults (Brooksbank et al., 1981; Coyle & Enna, 1976), as was done with the brain weight. We found two continuous measurements for this parameter for both species, one in the cortex and the other in the cerebellum. The cerebellar curve presents a hard to solve problem. The curve for humans presents a peak near birth while the rat does not show this pattern. In the graph it can be interpreted as a regression in development, which does not make theoretical sense. This is a limitation of this analysis that can only compare processes whose trends are in the same direction. Likewise, our model satisfactorily includes the curve within its margins (figure 3 supplementary material).

This problem was not observed in the case of GAD activity in the cerebral cortex. The last four graphs in Figure 5 show the activity of the enzyme for humans and rats throughout development. In this case, quantile regression also satisfactorily predicted the intervals within which the curve could be expected. However, GAD activity in the cerebral cortex was even less accurate than the brain weight to predict the distribution of developmental milestones.

**Figure 5:**
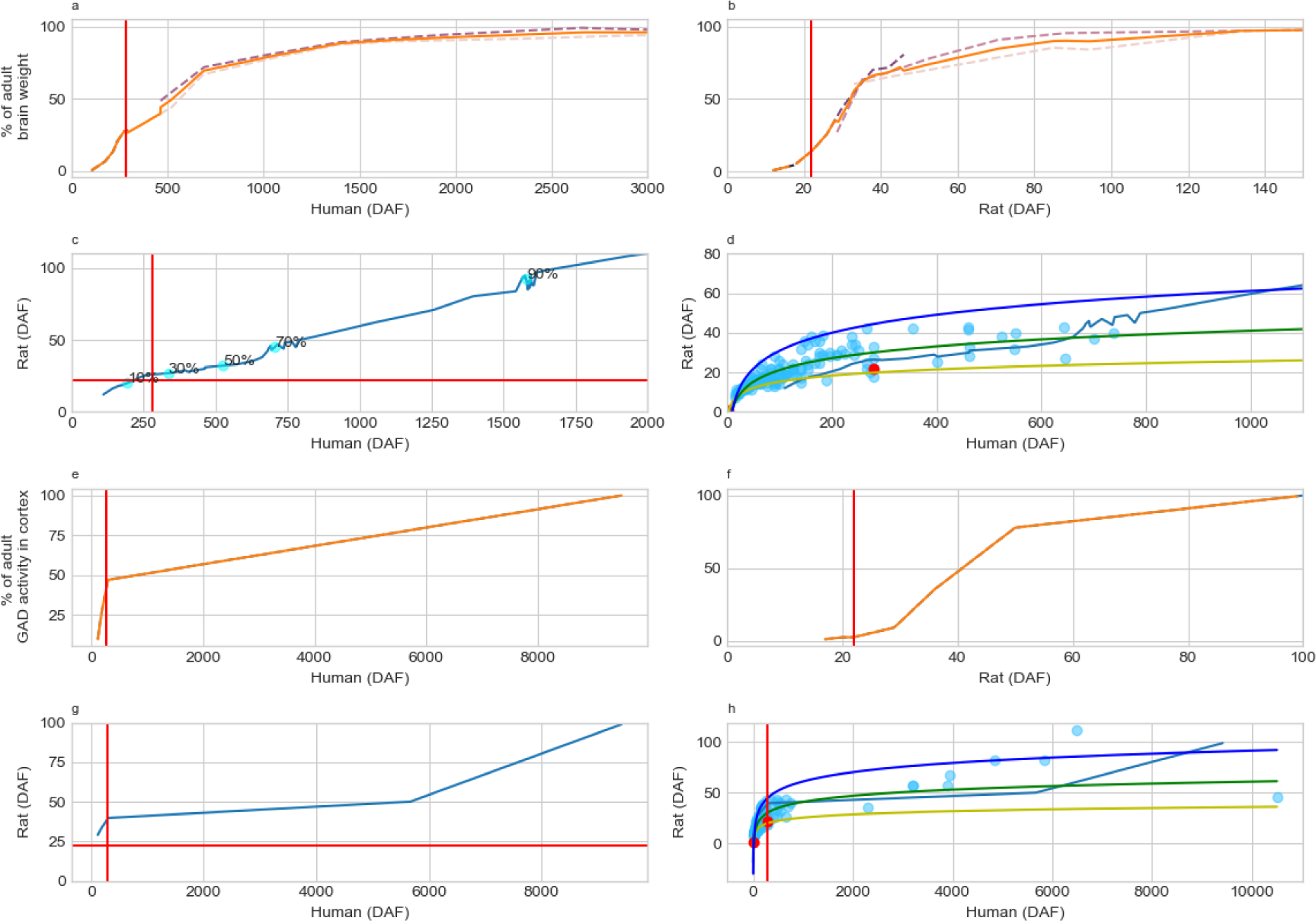
The first four subfigures plot aspects of comparative brain weight growth. The first two figures in the first row indicate the change in brain weight in percentage of the average adult value as dependent variable and days after fertilization (DAF) as independent variable for humans (a) and rats (b), respectively. The dotted lines indicate data from different publications, while the continuous line represents the average. In the first figure of the second row is plotted the isometric relationship between percentage of brain weight increase for both species, with some of these percentages also indicated on the plot (c). The second figure of the second row shows how these measurements overlap with our model of ranges of brain development (d).

## Discussion

With the intention of creating a mathematical model that explains the developmental relationships of rats and humans, a database with paired data of the development of humans and rats was generated. In addition, we combined our database with other databases from elsewhere. Previous works in this subject could not reach an agreement about the best model to compare the brain development of both species, and also failed to make a comprehensive and didactic interpretation of the models in order to be used by the scientific community.

### Clancy vs. Ohmura

After reading this work, there should be no doubts about the lack of linearity of the developmental relationships of both species. A logarithmic model fits better than a linear model. In this sense, we should agree with the model proposed by Clancy et al. (Clancy et al., 2001). Furthermore, we modified the proposed model to make it more specific to compare humans and rats. We also adjusted the parameters to translate postconceptional human days to postconceptional rat days, making the model easier to interpret. As criticism to Clancy’s model, it could be mentioned that some of their assumptions do not seem to be fulfilled. For example, the assumption that development processes follow the same order in humans than in other species. Although, there are developmental milestones that must necessarily occur concatenated and in order. For example, the cerebral cortex must be at least partially formed for GABAergic neurons to be able to migrate into it. There are also divergent processes where cell fates branch out and follow time patterns that are different from each other. In this case, the order in which the different development milestones occur could be altered. This is important because it changes the way development relationships should be interpreted. They should not be interpreted as a univocal relationship, but as relationships where a specific moment in the development of one species corresponds to a time range in the other. Secondly, the work of Clancy did not intend to be specific to compare rats and humans and include a small sample for the rat.

The work of Ohmura et al. (Ohmura & Kuniyoshi, 2017) has virtues and defects; its main virtue is having made the best database of development events to date, a relevant database even for the confection of this work. As criticisms, it should be noted that the proposal of the linear model is not very generalizable and only works for that period studied by the authors. The authors themselves clarify that if they included birth in the model, it ended up adjusting better to a logarithmic curve. Any nonlinear process can be fitted to a linear model at short ranges. The problem is that the model’s predictions outside the specific range are questionable, and any type of extrapolation loses precision. Science is a collective process, and the work of Ohmura and collaborators made an important contribution to general knowledge beyond the fact that their conclusions can be questioned.

### Methodological considerations and limitations

### Data science and biomedicine

Data science is revolutionizing both science and society. It has an interdisciplinary origin and continues providing mathematical and statistical tools to other areas of research. For example, quantile regression is not a new concept, it was proposed by Ruđer Josip Boskovic in 1760. However, it was recently incorporated into a library that allows its use with the python programming language. Relatively recently it was also developed for non-linear regressions (Geraci, 2019) but as far as we know it has not yet been incorporated into any python library.

### Challenges

Apart from the already mentioned fact that the data does not fit a linear model, other assumptions of ordinary least squares regressions are also not fulfilled. Developmental events are not independent of each other, for example the brain cannot grow in weight if the neural tube has not been formed yet. The data is not homoscedastic either since the variance increases with age. In this sense, the nonlinear regression by ordinary least squares can be used to determine the parameters of the model in an unbiased way but not to perform statistical inferences. For this reason, bootstrapping was used to obtain the 95% confidence intervals. When the common logarithm or the two-parameters logarithmic function (equation 1) were utilized to linearize the data there seems to be a change in slope around the time of birth that does not allow the data to be fully linearized. This is not a problem, on the contrary, it is an expected theoretical aspect of the relationship between the development of both species. The rat can have 16 pups in a litter while humans invest all that energy usually in a single offspring. The alternative is to separate the model in 2 but we prefer to apply the principle of parsimony and maintain a single model. The least-squares curve is similar to the mid-quantile curve, with only about a 10% difference at 27 years of human age.

With respect to the contiguous data, the curves sometimes present strange aspects and edges (Figure 5). This occurs because, we decide to perform linear interpolations, the sum of linear interpolations sometimes ends up giving that saw-like appearance. It should not generate important errors, at least within the intervals that comprise 90% of the data. Outside these intervals, a small change in one of the axes can generate a large change in the other axis. This problem has already been described by Clancy (Clancy 2001). We agree on their evaluation and applied the same solution, which is to discard the most external data. Linear interpolations have some advantages, the main one being their simplicity, but they also have the advantage that they do not hide data series with few values as in the case of GAD activity data.

### Final considerations

In this work, a public database was generated that can be used to analyze the relationship of development between humans and rats. Also, a mathematical model was proposed to choose the best age for a rat model to resemble a particular period of human brain development. The aim of this work was to make the model as simple as possible, didactic, easy to understand and feasible to use. Both the program and the database are available for consultation and for other researchers to carry out their own analyses. Widely accepted concepts in evolution can be applied to interpret the developmental relationship proposed in this work. First, the development of animals of different species is more similar the earlier the event is observed, second larger animals usually need more time to develop, see (Abzhanov, 2013; Richardson, 2022) for more insight about these topics. Of course, there are multiple factors that make these rules more complex, but a non-linear development relationship is clearly to be expected.

## Supporting information

supplementary files statsnl.txt

statsolslr.txt

## Notes

https://vazquez-borsetti.github.io/rat-and-human-comparative-development/

### Competing Interest Statement

The authors have declared no competing interest.

