## supplementary figure for "Divergent pattern of development in rats and humans"

Supplementary figures 1 and 2.

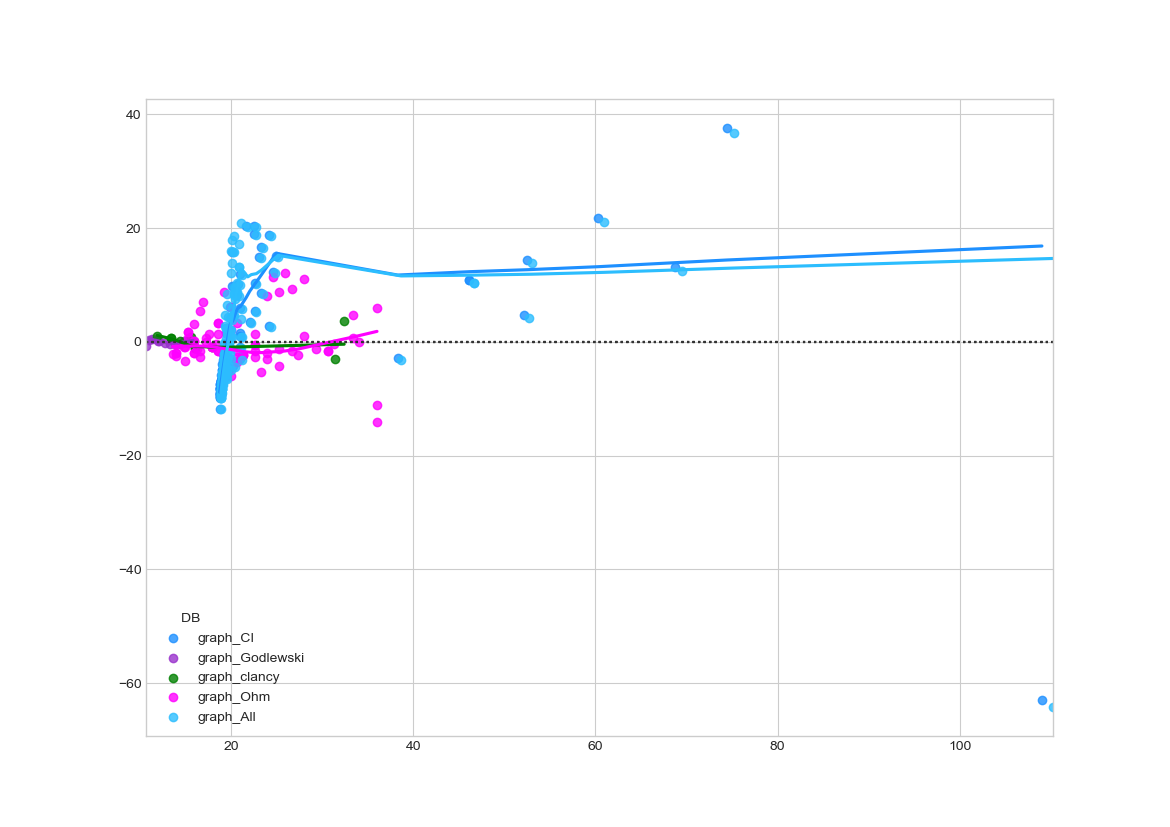

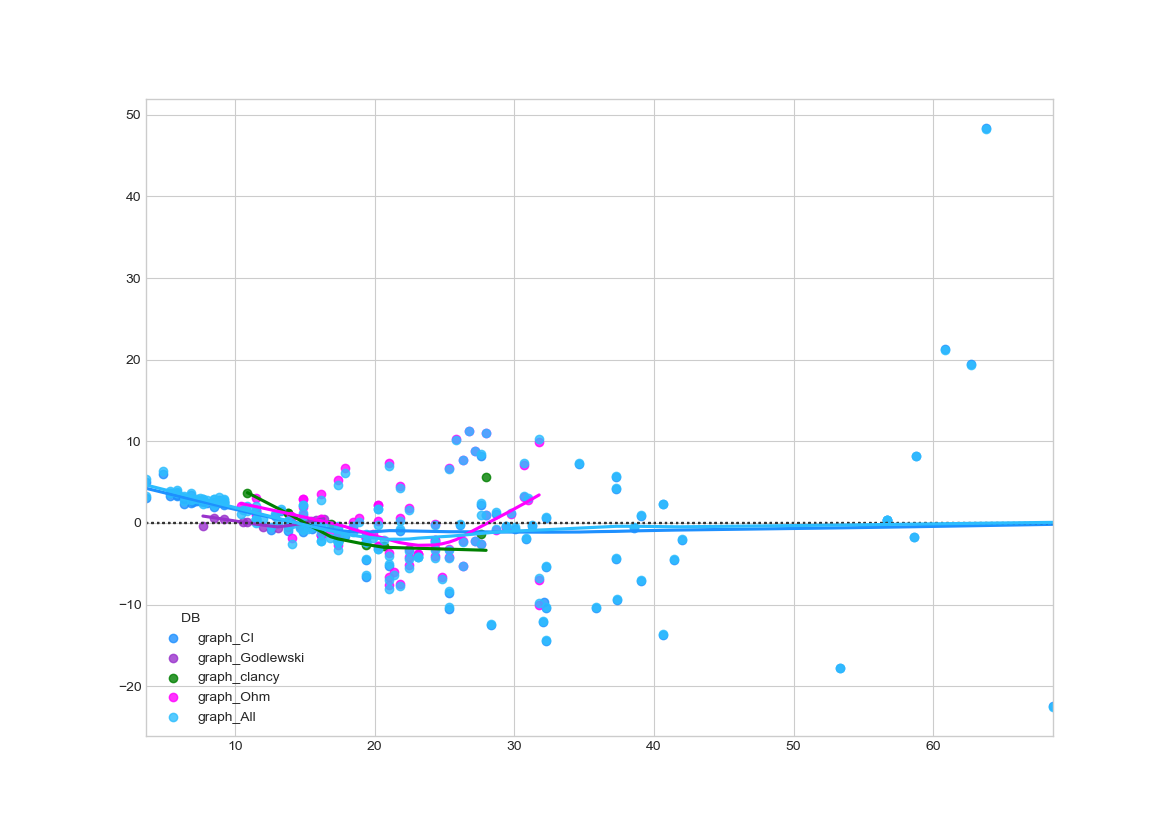

In these figures are displayed the residuals of both models. The supplementary figure 1 corresponds to the linear regression. It is evident that this kind of regression cannot explain late developmental events and is not consistent among different databases. On the other hand, the logarithmic model can describe all the data and is consistent among different databases. In the logarithmic model, can be also spotted a putative point of inflection at birth.

supplemental image 3.

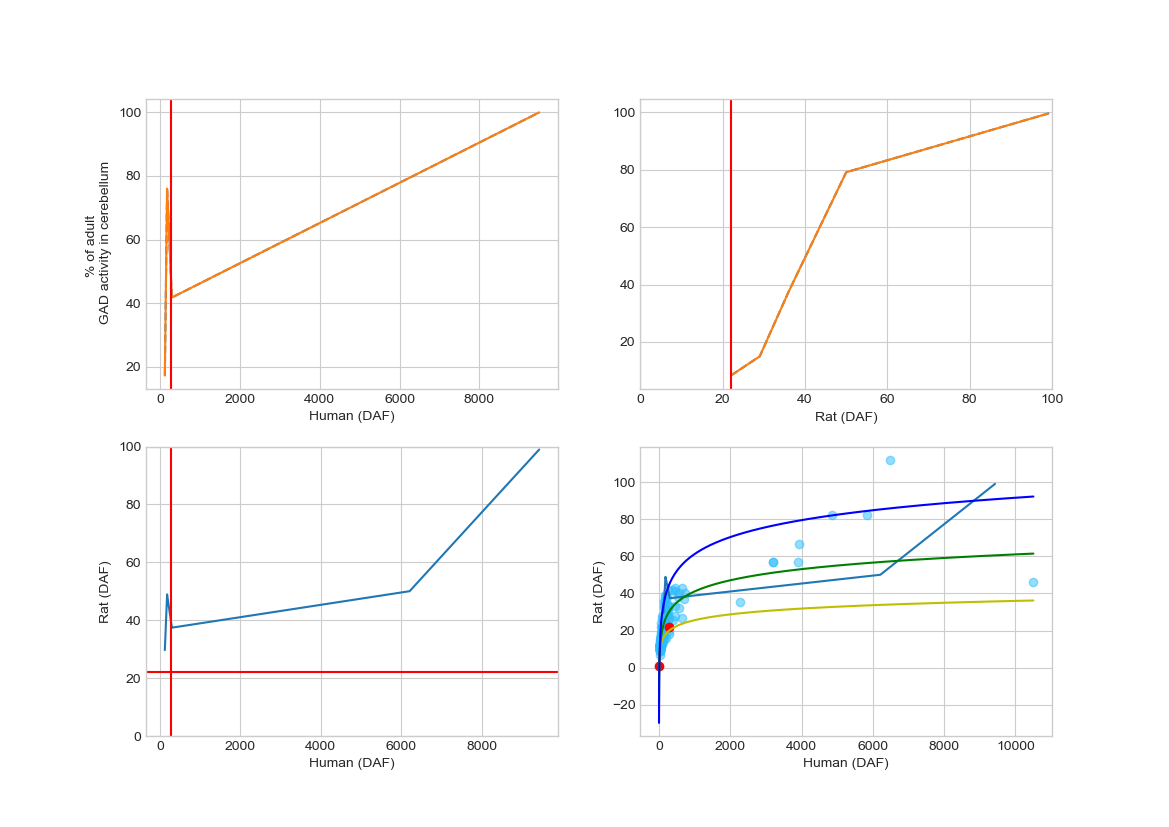

Like figure 5 here is plotted quantitative continuous data on GAD enzyme activity, in this case in the cerebellum. The two graphs in the first row show the increase in enzyme activity, as percentage of the average adult value on days after fertilization, throughout development for humans and rats, respectively. The figures in the last row show the isometric relationship for the activity of the enzyme.

Supplemental image 5 and 6.
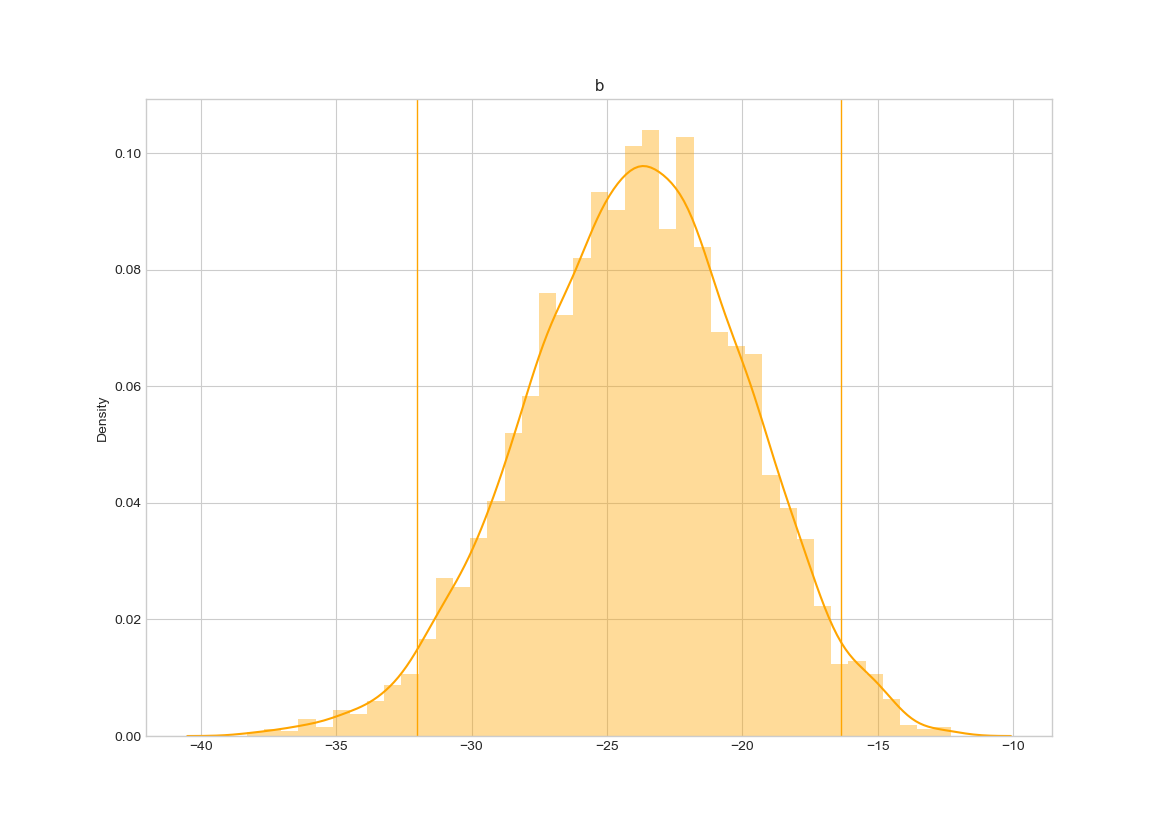

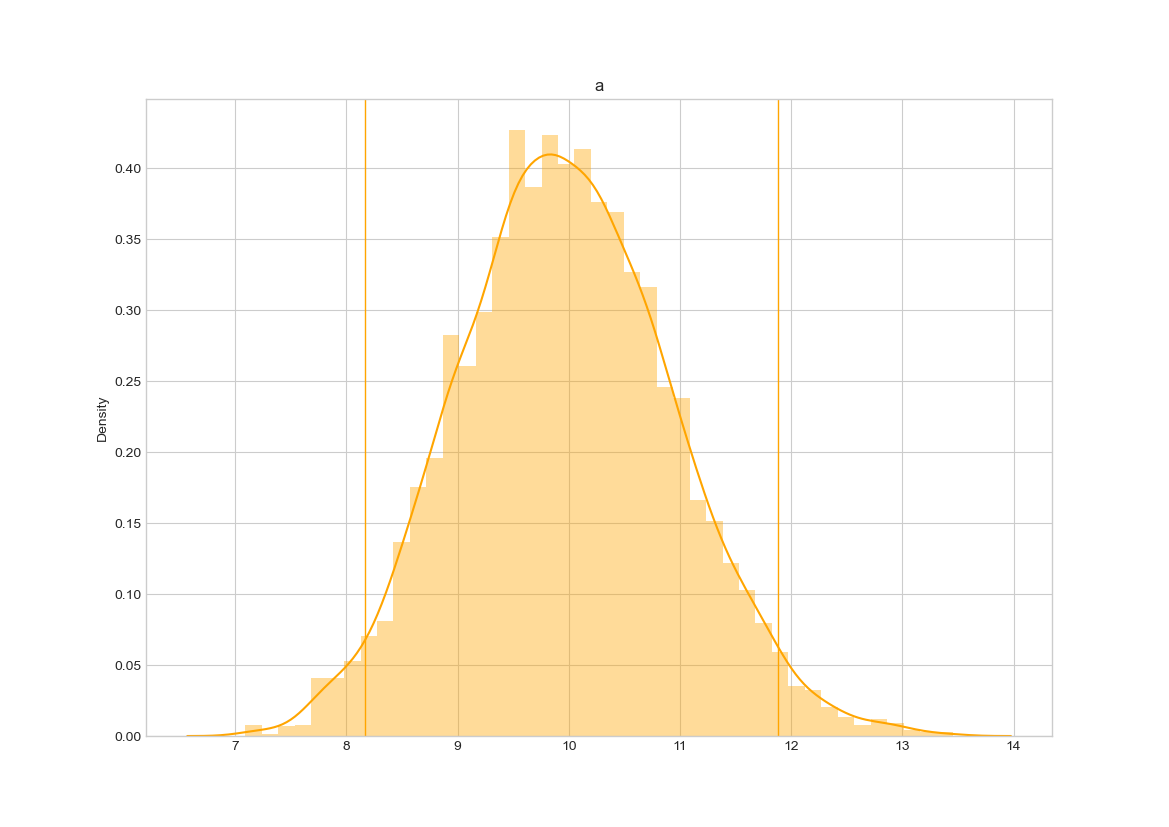

The bootstrapping of the parameter a and b of the nonlinear regression (equation 1)

supplemental image 7.

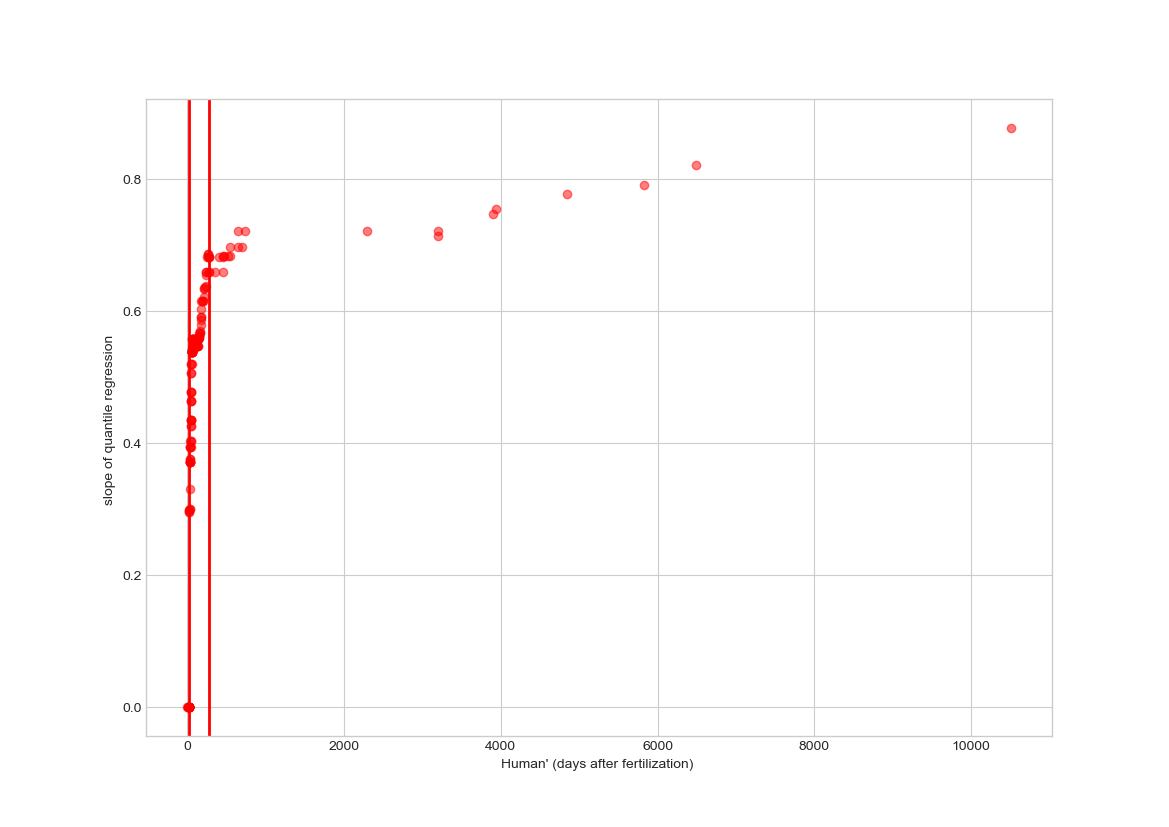

The slope of the median quantile as a function of the human age. The inflection point at birth (second vertical line) becomes evident.
